# The association of Mediterranean plant species with herbivorous arthropods and its effect on pest abundance in organic vineyards

**DOI:** 10.1101/2024.11.12.623178

**Authors:** Renata Santos, Pedro Naves, Rita Morais, Leonor R. Rodrigues, Sílvia Pina, Márcia Santos, Maria Teresa Rebelo, Patrícia Garcia Pereira, Sara Magalhães, Elisabete Figueiredo

## Abstract

Non-crop plant resources, such as hedgerows and adjacent woodland areas, may impact the distribution of pest species in the crop. Knowledge on the associations between plants and arthropods as well as their impact on pest distribution is thus key to adequately manage agroecosystems. We selected a number of native Mediterranean plant species located around organic vineyards in Southern Portugal and determined their associations with species of Auchenorrhyncha and phytophagous tetranychid and tenuipalpid mites, including the main species of vineyard pests in the area. We also tested if the abundance of vineyard pests is affected by the distance to the edge and the species of plants present. Most non-crop plants and ground cover vegetation harbored very low numbers of leafhopper pests. *Rubus ulmifolius* and *Tamarix africana* proved to be a repository of non-pest Auchenorrhyncha species, with the former also serving as winter repository of the pest *Jacobiasca lybica. Rosa canina* and *Fraxinus angustifolia* hosted abundant populations of the spider mite *Tetranychus urticae*. Still, plots next to plant biodiverse margins harbored fewer numbers of *T. urticae*, when compared with plots next to other vineyards. Furthermore, pest abundance in vineyards increased with growing distance to plant biodiverse margins. Our results highlight the benefits of biodiverse margins in reducing pest abundance and point to the importance of a good selection of plant species when managing and planning these non-crop plant resources.

## Introduction

Native to the Mediterranean basin, the common grapevine (*Vitis vinifera* L., Vitaceae) is an important and emblematic crop in all regions with a Mediterranean climate. However, its continued expansion and intensification can be a cause of habitat destruction and loss of biodiversity (Viers et al. 2013). Moreover, the simplification of complex vineyard landscapes can lead to a simultaneous reduction of insect biodiversity and increase in pest abundance (Altieri and Nicholls 2002). Therefore, there is a need to adopt strategies and management practices that make viticulture more sustainable (Viers et al. 2013). Although organic production is widely seen as an alternative with less impact on biodiversity than conventional farming (Tuck et al. 2014), its environmental sustainability remains an ongoing debate (Froidevaux et al. 2017; Meemken and Qaim 2018).

One of the main challenges of organic management is the implementation of effective crop protection measures (Meemken and Qaim 2018). These practices include the creation and maintenance of non-crop plant resources, such as hedgerows and woodland areas in the margins of crops. By providing resources and refuges for natural enemies throughout the year, these areas can positively impact biological control of pest species inside crops (Begg et al. 2017; Holland 2019; Paredes et al. 2019), including vineyards (Altieri and Nicholls 2019; Favor et al. 2024; Gavinelli et al. 2020; Thomson and Hoffmann 2009, 2013). However, non-crop plants can also serve as habitat and food sources for pests and thus fail to enhance pest control (de Villiers and Pringle 2011; Holland 2019; Lu et al. 2014; Morandin et al. 2011; Ponti et al. 2005; Tscharntke et al. 2016; Villa et al. 2020; Wilson et al. 2016). The role of such marginal areas in crop management hinges upon the associations between species of plants and arthropods. It is thus key to select the best combination of plants to include in ecological infrastructures (Altieri and Nicholls 2019; Duso et al. 2012; Holland 2019; Morandin et al. 2011). Therefore, to adequately manage agroecosystems, it is key to locally assess the impact of marginal areas in the abundance of crop pests.

In Portugal, several arthropod groups cause great damage to vineyards. The most common ones are highly polyphagous vineyard pests common in the Mediterranean basin: leafhoppers, such as the cotton jassid, *Jacobiasca lybica* (Bergevin & Zanon) (Hemiptera: Cicadellidae) (Lopes et al. 2014; Quartau and Rebelo 1992; Tsolakis & Ragusa 2008), and the two-spotted spider mite, *Tetranychus urticae* Koch (Trombidiformes: Tetranychidae) (Duso et al. 2012; Naves et al. 2021; Tsolakis and Ragusa 2008). Of phytosanitary importance are also several species of froghoppers, spittlebugs and leafhoppers that are, in Europe, potential vectors of *Xylella fastidiosa* (Wells et al.), a plant pathogen which can affect several cultivated and wild plants (Di Serio et al. 2019; EFSA PLH Panel 2015; Lopes et al. 2014; Moussa et al. 2016; Villa et al. 2020). In Mediterranean regions, vineyard pests, such as leafhoppers and spider mites, have been shown to cyclically move between the crop and non-crop host plants (Duso et al. 2012; Wilson et al. 2016). Likewise, specific non-crop plant species can be beneficial to spider mites’ natural enemies, such as Phytoseiidae mites, and leafhopper natural enemies, such as *Anagrus* sp. parasitoids, which disperse from surrounding natural areas to the crops (Duso et al. 2004, 2012; Gavinelli et al. 2020; Ponti et al. 2005; Wilson et al. 2016). However, even if positive, the impact of non-crop plant species tends to decrease with increased distance to these peripheral natural resources. Therefore, opposite gradients of decreased biological control and increased pest abundance with increased distance to these peripheral resources are expected and have been observed in vineyards, in leafhoppers (Altieri and Nicholls 2019) and other pests (Thomson and Hoffmann 2013). Hence, marginal vegetation may either positively or negatively affect hemipteran and tetranychid pests. It is thus important to properly identify the non-crop plant species that harbor such crop pests as well as the specific impact of marginal plants on pest abundance and distribution in vineyards.

Here, we set out to address these issues. As in the Mediterranean basin herbaceous ground covers tend to dry out completely in late spring and summer, we focused on perennial plant species present in hedgerows and peripheral areas around vineyards, which remain green for the entire dry season. These plants also tend to be less well studied than the herbaceous layer (Holland 2019). We thus selected a number of common spontaneous or planted native Mediterranean plant species to identify which act as pest repositories, or as hosts of other Auchenorrhyncha and phytophagous mites that may be alternative prey to natural enemies or potential *X. fastidiosa* vectors. We also assessed whether the distance to the margins affected the abundance of such crop pests in the vineyard. This study contributes to evaluate the role of different plant species at the crop margin in pest abundance and distribution, contributing to more effective strategies of agroecological practices in vineyards.

## Materials and methods

### Study site

Field trials were conducted at Herdade do Esporão (38°22’48.23”N, 7°33’39.52”W, 206 m altitude) in interior Alentejo, southern Portugal (Fig. 1). This is a 1748 ha estate comprising Mediterranean open forests (*montado* agro-systems) dominated by holm oak (*Quercus rotundifolia* Lam.), shrublands, riparian forests, 441.5 ha of vineyards and 93.3 ha of olive groves, fully converted to certified organic farming since 2017. The dominant soils are derived from schist, diorite and granite and the climate is a typical Southern Mediterranean climate, fitting within the Csa Koppen-Geiger climate classification (AEMET 2011).

**Fig. 1.**
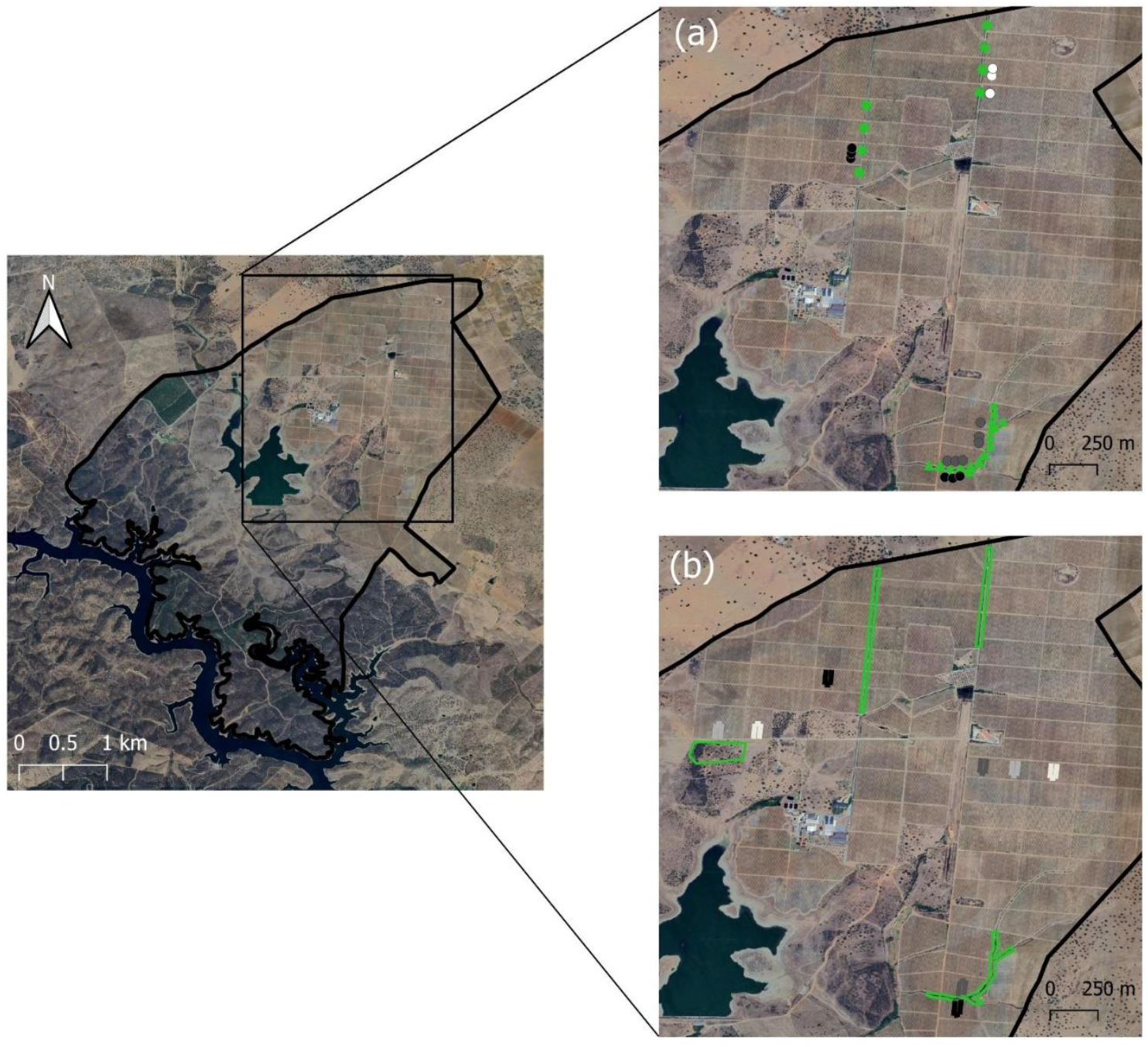
Location of the study area in Alentejo, southern Portugal: A) location of the transects for Auchenorrhyncha sampling: ‘Alicante Bouschet’ vineyards - black balls; ‘Syrah’ vineyards – grey balls; ‘Aragonez’ vineyards – white balls; planted hedgerows – green asterisks; Rosaceae-dominated hedgerows – green triangles; *Tamarix*-dominated hedgerows – green squares; B) location of phytophagous mites’ sampling areas: (blue – ‘Alicante Bouschet’ vineyard plots – black, ‘Syrah’ vineyard plots – dark grey; ‘Cabernet Sauvignon’ vineyard plots – light grey; ‘Touriga Franca’ vineyard plots – white; marginal areas are in green.

### Auchenorrhyncha sampling design

Twenty-four 25 m transects were established along three types of hedgerows bordering different vine cultivars: eight transects in two parallel planted hedgerows (PH) located along a road (separated from each other by a minimum of 125 m); eight transects in spontaneous hedgerows dominated by Rosaceae (RH) located along an irrigation ditch (separated from each other by 30 m); and eight transects in spontaneous hedgerows dominated by *Tamarix africana* Poir. (TH) also located along an irrigation ditch (separated from each other by 30 m) (Fig. 1a). All hedgerow corridors were connected to areas with holm oak *montado* (HOM). Transects were fixed and a maximum of two plants per species per transect was randomly selected and marked for sampling.

Between 2021 and 2023, on six dates - in spring (May 2021 and 2022), summer (early September 2021 and August 2022) and winter (January 2022 and 2023) - eight native tree and shrub species were sampled (Table 1). Marked plants were sampled on each date, apart from *Rosa canina* L. and plants from TH which were not sampled in January. Not all plant species were present in all transects in each hedgerow. *Nerium oleander* L. was the only plant present in all hedgerow types (PH, RH and TH). Spontaneous herbaceous ground cover plants were also recorded and sampled once per transect. Standardized 30 s suction samples were taken from the selected plants (from the stem to the outermost leaves) and from the ground cover (while walking along the transect at a standard pace) using a STIHL SH 86 C-E garden vacuum adapted for invertebrate sampling with the insertion of an anti-trip mesh on the vacuum tip.

**Table 1.**
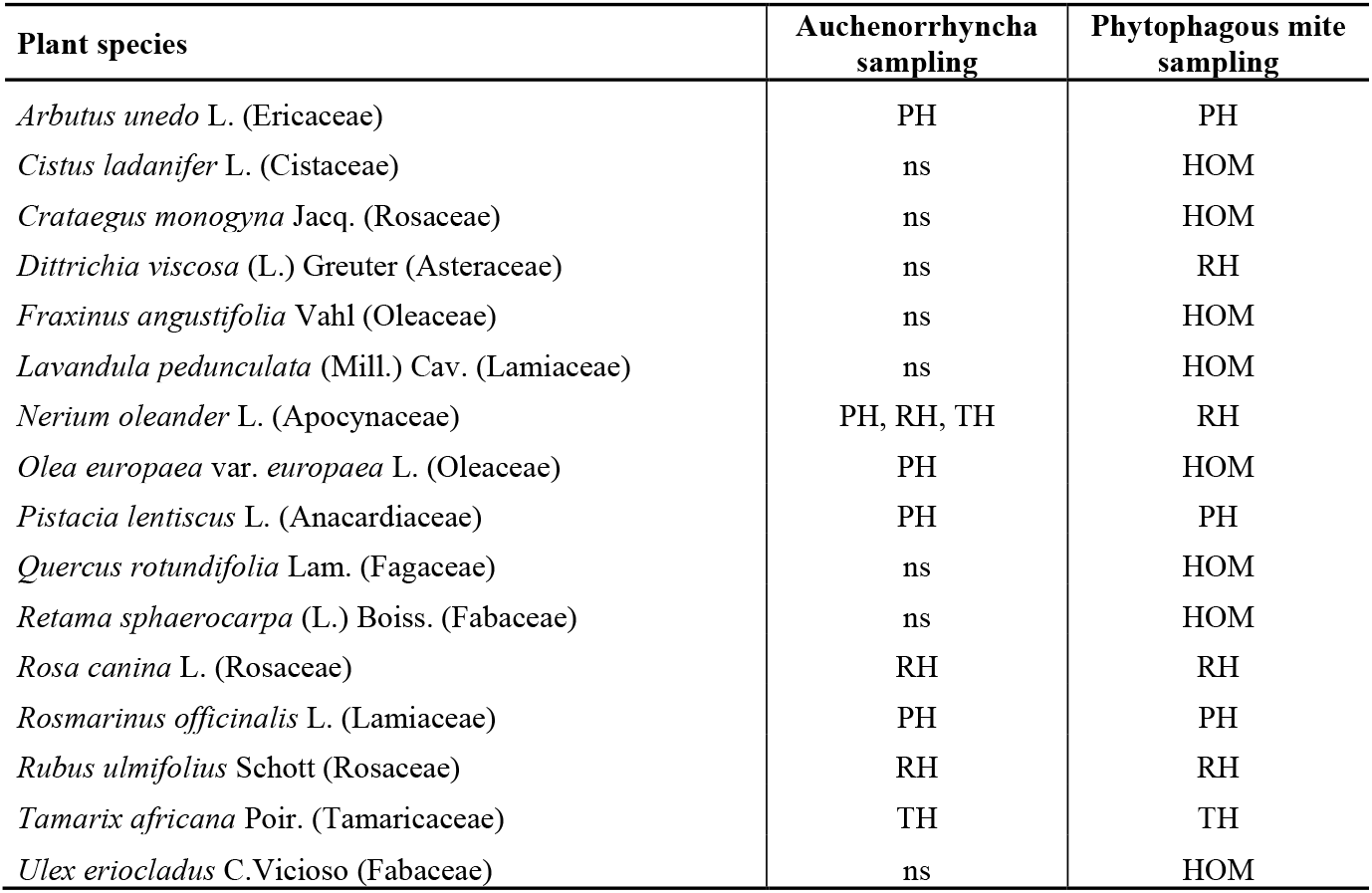
Plant species sampled for Auchenorrhyncha (by suction), and for phytophagous Tetranychidae and Tenuipalpidae mites (by leaf collection), by habitat type. HOD – Holm oak *montado*; PH – Planted Hedgerows; RH – Rosaceae-dominated Hedgerows; TH – *Tamarix*-dominated Hedgerows; ns – not sampled.

Another 15 transects were established, perpendicularly to the sampled hedgerows, on adjacent vineyards of three red cultivars: in ‘Alicante Bouschet’ three transects perpendicular to PH and three perpendicular to RH; in ‘Syrah’ three transects perpendicular to RH and three perpendicular to TH; and in ‘Aragonez’ three transects perpendicular to PH (Fig. 1a). Transects were at a minimum distance of 40 m from each other. In May and August, in each transect, two to four vines were sampled (30 s suctions) at three distances from the hedgerows: 10 m, 50 m and 100 m. ‘Alicante Bouschet’ vineyards were sampled in 2021 and 2022, other casts only in 2022. Sampling was carried out between 9 am and 18 pm, in mild wind (below 4 in the Beaufort Wind Scale (WMO 1970)) and mild to high temperatures (9 °C to 35 °C), taken from the local weather forecast. Each sample was stored in 70% ethanol. All samples were observed under a stereoscopic microscope and Auchenorrhyncha specimens were sorted and counted.

In July 2020 and 2022, each of the ‘Alicante Bouschet’ vineyards next to PH and RH were surveyed for leafhopper attack levels, by counting the number of Empoascini (*J. lybica* tribe) nymphs in 100 leaves of the eastern side of 50 random vines (2 leaves/vine: 7^th^ and 8^th^ leaf). Economic threshold is reached when 50 or more nymphs are counted per 100 leaves (DRAPA 2018).

### Phytophagous mites sampling design

In July and September 2021 and in July 2022, 50 to 150 leaves were collected from five individual plants (10 to 30 leaves per plant) of sixteen native Mediterranean species, found in different habitats next to vineyards (Fig. 1b; Table 1).

In July 2021 and 2022, grapevine leaves were collected from four red cultivars: ‘Alicante Bouschet’, ‘Syrah’, ‘Cabernet Sauvignon’ and ‘Touriga Franca’. In each cultivar, two plots of 30 m x 100 m were delimited: one next to a pathway surrounded by other vineyards (central plot) and another next to a biodiverse periphery composed of hedgerows or holm oak *montado* (peripheral plot) (Fig. 1b). In each selected plot, 60 grapevine leaves were collected, comprising 30 samples: 2 leaves/plant (one sample), in 10 consecutive lines, at 20 m, 60 m and 100 m from the periphery (10 samples per distance). All leaves were kept in plastic bags at 5 °C until they were observed under a stereoscopic microscope and mites retrieved manually to 70% ethanol or AGA solution (ethanol, glycerol and acetic acid) (Gutierrez 1985).

In July 2020 and 2022, all plots were surveyed for spider mite attack levels, by checking the presence of spider mites in 50 random vines per plot (2 leaves/vine: from the middle of the shoot). Economic threshold is reached when at least 30-45% of leaves are occupied (DRAPA 2018).

### Species identification and data analysis

Auchenorrhyncha specimens were identified to the lowest possible taxonomic level using identification keys and with the help of experts. In most cases, male genitalia are essential for identification and only males can be identified to the species level. All individuals belonging to the family Aphrophoridae plus the ones belonging to the species *Euscelis lineolata* Brullé (Cicadellidae) were considered to be potential *X. fastidiosa* vectors (EFSA PLH Panel 2015; Moussa et al. 2016).

Phytophagous mites were mounted and prepared for identification. Specimens were cleared in lactic acid (50 %) for a day and mounted in Hoyer’s medium for phase contrast microscope observation. Identification was based on morphological characters of both sexes, using available descriptions.

Either Generalized Linear Models (GLMs) or Generalized Linear Mixed Models (GLMMs) were fitted, assuming Poisson (function *glm*, stats package and function *glmer*, lme4 package, respectively; Bates et al. 2015; R Core Team 2023) and Negative Binomial error distributions (function *glm*.*nb*, MASS package and function *glmer*.*nb*, lme4 package, respectively; Bates et al. 2015; Venables and Ripley 2002), tested for overdispersion and selected based on Akaike information criterion (function *AICtab*, bbmle package; Bolker and R Development Core Team 2023). The statistical significance of fixed effects was estimated based on Chi-squared (χ^2^) tests (Anova, *car* package; Fox and Weisberg 2019). To simplify maximal models and establish a minimal model, non-significant terms were sequentially eliminated from the highest to the simplest order interaction (Crawley 2007). Random effects were removed when they did not significantly improve the model fit. To interpret significant effects (*p* < 0.05) of factors with three or more levels and their interactions, multiple comparisons were performed (pairs, *emmeans* package; Lenth 2023). All tests were performed with RStudio software version 2023.06.1+524 (R Core Team 2023).

The effect of plant species, month and year (fixed factors) on the abundance of Empoascini and other Auchenorrhyncha at field margins was tested using Poisson and Negative Binomial error distributions, respectively, with transect and habitat as random factors (*J. lybica* alone was not tested due to very low numbers). To assess associations among Auchenorrhyncha and plant species, a model was constructed with abundance of males of common species (global abundance of N≥1% of individuals) as the response variable, plant species as fixed factor, and hedgerow type, transect, month and year as random factors. *N. oleander* from different types of hedgerows (PH, RH and TH) were tested as different species.

The effect of hedgerow type, month and year (fixed factors) on the abundance of total Auchenorrhyncha on the ground cover was tested using Negative Binomial error distribution (Empoascini was not tested due to very low numbers). To assess associations among Auchenorrhyncha species and ground cover of the different types of hedgerows, a model was constructed with abundance of males of common species as the response variable, hedgerow type as fixed factor, and month and year as random factors.

In the vineyard, to test how distance to the periphery affects the abundance of *J. lybica* in August, four models were run: a) a model including all ‘Alicante Bouschet’ samples from 2021 and 2022, with distance to the periphery as covariate, and plot and year as fixed factors, b) a model including samples of all cultivars of 2022, with distance to the periphery as covariate, and cultivar as fixed factor and c) two models including either ‘Alicante Bouschet’ or ‘Syrah’ samples from 2022, with distance to the periphery as covariate and type of periphery as fixed factor.

To test how distance to the periphery affects the abundance of *T. urticae*, we ran a model including all samples, with distance to the periphery as covariate, and plot type (peripheral/central), cultivar and year as fixed factors. Upon reaching multiple significant triple interactions, the data was first subset into the two years, and then subset once more by cultivar, in order to look at more general abundance patterns within each year and cultivar specific effects, respectively. In both new models, distance to the periphery was set as covariate and plot type as fixed factor.

## Results

### Auchenorrhyncha

In the hedgerow’s perennial plants, a total of 3911 Auchenorrhyncha, belonging to at least 42 species, were collected (Table S1). Of these, 208 (5.3%) were Empoascini and, of the males identified to species level, nine were *J. lybica*, 50 *Hebata solani* (Curtis), and two *Hebata decipiens* (Paoli) (*Hebata* is a reclassification of the genus *Empoasca* by Xu et al. 2021, a change widely accepted by other Auchenorrhyncha specialists: Dmitriev et al. 2024; Evangelou et al. 2023; Nickel 2022). Males of *J. lybica* were collected in *R. ulmifolius* (seven in January and one in August) and *A. unedo* (one in January). Overall, only 26 specimens (0.7%) belonged to species considered potential vectors of *X. fastidiosa* (Table S1).

Plant species (145.662, *df* = 9, *p* < 0.001), month (*χ*^2^ = 25.980, *df* = 2, *p* < 0.001), year (*χ*^2^ = 5.063, *df* = 1, *p* < 0.05) and the interaction between plant species and month (*χ*^2^ = 57.631, *df* = 15, *p* < 0.001) significantly affected the abundance of Empoascini, but interactions involving year did not (Table S2). Similarly, plant species (*χ*^2^ = 1251.71, *df* = 9, *p* < 0.001), month (*χ*^2^ = 30.91, *df* = 2, *p* < 0.001) and the interaction between plant species and month (*χ*^2^ = 45.00, *df* = 15, *p* < 0.001) significantly affected the abundance of other Auchenorrhyncha, but year and its interactions did not (Table S2). Specifically, *R. ulmifolius* and *R. officinalis* harbored higher numbers of Empoascini in January, with the latter also harboring high numbers in May. In August, all plants had very low numbers of Empoascini. Regarding other Auchenorrhyncha, higher numbers were harbored by *T. africana* in May and August, and by *R. ulmifolius* in January (Fig. 2). The abundance of Empoascini only varied between months in *R. officinalis* and *R. ulmifolius*. For other Auchenorrhyncha, significant abundance variations between months also occurred in *A. unedo, P. lentiscus* and *T. africana* (Table S3).

**Fig. 2.**
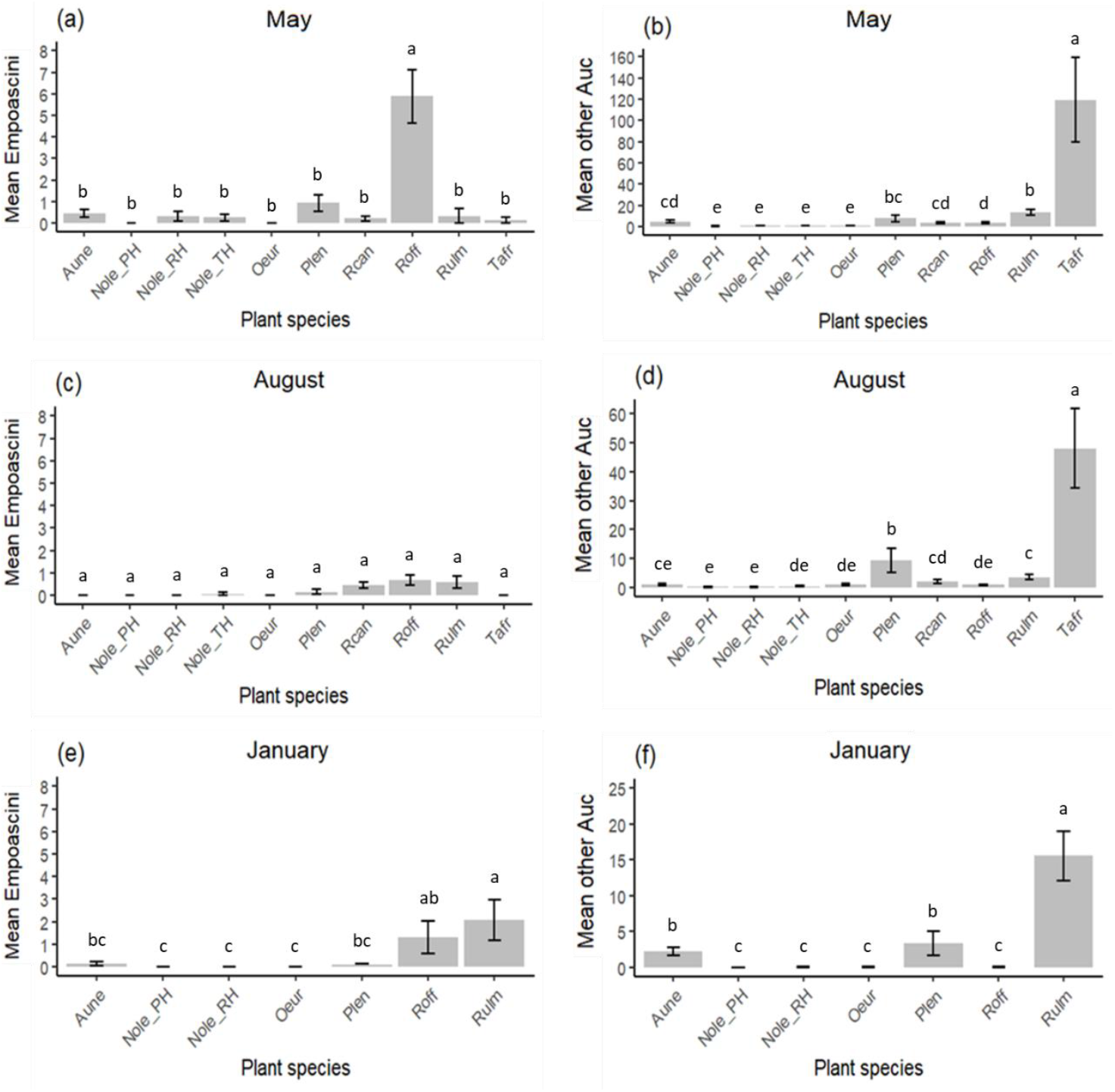
Mean abundance/sample ± standard error per plant species of a) Empoascini and b) other Auchenorrhyncha (Auc) in May; c) Empoascini and d) other Auchenorrhyncha (Auc) in August; e) Empoascini and f) other Auchenorrhyncha (Auc) in January. Letters above bars denote significance of multiple comparison tests at α = 0.05. ‘Aune’ - *Arbutus unedo*; ‘Nole_PH’ - *Nerium oleander* in Planted Hedgerows; ‘Nole_RH’ - *Nerium oleander* in Rosaceae-dominated Hedgerows; ‘Nole_TH’ - *Nerium oleander* in *Tamarix*-dominated Hedgerows; ‘Oeur’ - *Olea europaea*; ‘Plen’ - *Pistacia lentiscus*; ‘Rcan’ - *Rosa canina*; ‘Roff’ - *Rosmarinus officinalis*; ‘Rulm’ - *Rubus ulmifolius*; ‘Tafr’ - *Tamarix africana*. Note the differences in scales.

Significant associations were detected between males of five common Auchenorrhyncha species and three plant species (Table 2). For *H. solani*, although GLM result showed only a marginally significant difference between plant species, multiple comparisons showed that *R. officinalis* harbors either significantly or marginally significantly more of this species than other plants (Table S4).

**Table 2.** Generalized linear model output for the association between perennial plant species and abundance of males of common Auchenorrhyncha species (global abundance of N≥1% of individuals) (results from multiple comparisons are shown in Table S4). *χ*^2^: chi-square value. *df*: degrees of freedom. ‘Plant species’: plants in all hedgerows (*Arbutus unedo, Nerium oleander* - tested separately for each hedgerow -, *Olea europaea, Pistacia lentiscus, Rosa canina, Rosmarinus officinalis, Rubus ulmifolius, Tamarix africana*). ‘var’ – variable.

| Variable of interest | Explanatory var. | $\chi^2$ | df | p value | Plant association |
| --- | --- | --- | --- | --- | --- |
| *Arboridia* sp. 1 | Plant species | 59.678 | 9 | < 0.001 | *Rubus ulmifolius* |
| *Bugraia ocularis* (Mulsant & Rey) | Plant species | 80.302 | 9 | < 0.001 | *Pistacia lentiscus* |
| *Duilius* sp. 1 | Plant species | 428.65 | 9 | < 0.001 | *Tamarix africana* |
| *Hebata solani* (Curtis) | Plant species | 16.338 | 9 | 0.060 | - |
| *Opsius stactogalus* Fieber | Plant species | 165.19 | 9 | < 0.001 | *Tamarix africana* |
| *Ribautiana tenerrima* (Herrich-Schäffer) | Plant species | 32.278 | 9 | < 0.001 | *Rubus ulmifolius* |

In the hedgerow herbaceous ground cover plants, a total of 1326 Auchenorrhyncha, belonging to at least 31 species, were collected. Of these, only 19 (1.4%) were Empoascini, corresponding to seven males of *H. solani* and one of *H. decipiens*. No *J. lybica* males were collected. Only 29 specimens (2.2%) were potential *X. fastidiosa* vectors (Table S1). When comparing abundance of total Auchenorrhyncha, hedgerow type proved non-significant (*χ*^2^ = 4.297, *df* = 2, *p* = 0.117), but month (*χ*^2^ = 84.807, *df* = 2, *p* < 0.001), year (*χ*^2^ = 7.433, *df* = 1, *p* < 0.01) and the interaction between hedgerow and month (*χ*^2^ = 55.972, *df* = 3, *p* < 0.001) proved significant (Table S5). Indeed, in August PH showed a significantly higher abundance of Auchenorrhyncha than RH, but in January the opposite was observed. In RH and TH Auchenorrhyncha abundance changed significantly between sampled months, but in PH it did not (Table S6).

Only one common Auchenorrhyncha species was significantly associated with the ground cover of one hedgerow type: *Duilius* sp.1 with TH (*χ*^2^ = 6.566, *df* = 2, *p* < 0.05) (Tables S7 and S8). But as seen above, this species showed a much stronger association with the plant *T. africana* and was probably sampled on the ground cover plants of TH due to the presence of low height *T. africana*.

In the vineyard, a total of 4070 Auchenorrhyncha, belonging to at least 22 species, were collected. Of these, 2864 (70.4%) were Empoascini (Table S9). Only 185 (6.5%) were collected in May, and here we found 13% *J. lybica* males, 80% of *H. solani* males and 7% *H. decipiens* males. Of the 2679 (93.5%) Empoascini collected in August, all males were identified as *J. lybica*, which confirms observations in previous years (Alvarez 2020). Only 64 (1.6%) specimens were potential *X. fastidiosa* vectors (Table S9).

In ‘Alicante Bouschet’ in August 2021 and 2022, the GLM confirmed that distance to the periphery (*χ*^2^ = 9.435, *df* = 1, *p* < 0.01), plot (*χ*^2^ = 265.944, *df* = 1, *p* < 0.001) and year (*χ*^2^ = 26.154, *df* = 1, *p* < 0.001) all significantly affected *J. lybica* abundance, with no significant interactions between these variables (Table S10). This means that *J. lybica* abundance increased significantly with increased distance to the periphery in both vineyards and years, but more of these insects were collected in 2021 than in 2022 and more specimens were found next to RH than next to PH (similar to Fig 4a). This is in accordance with visual surveys, which showed that the vineyard next to RH reached economic threshold levels for leafhopper attack in both 2020 and 2022, while the one next to PH did not.

For data collected in all cultivars in August 2022, the distance to the periphery (*χ*^2^ = 5.121, *df* = 1, *p* < 0.05) and cultivar (*χ*^2^ = 78.042, *df* = 2, *p* < 0.001) significantly affected *J. lybica* abundance, but their interaction did not (*χ*^2^ = 0.553, *df* = 2, *p* = 0.758). Therefore, *J. lybica* abundance increased with increased distance to the periphery (hedgerows) in all cultivars (Fig. 3).

**Fig. 3.**
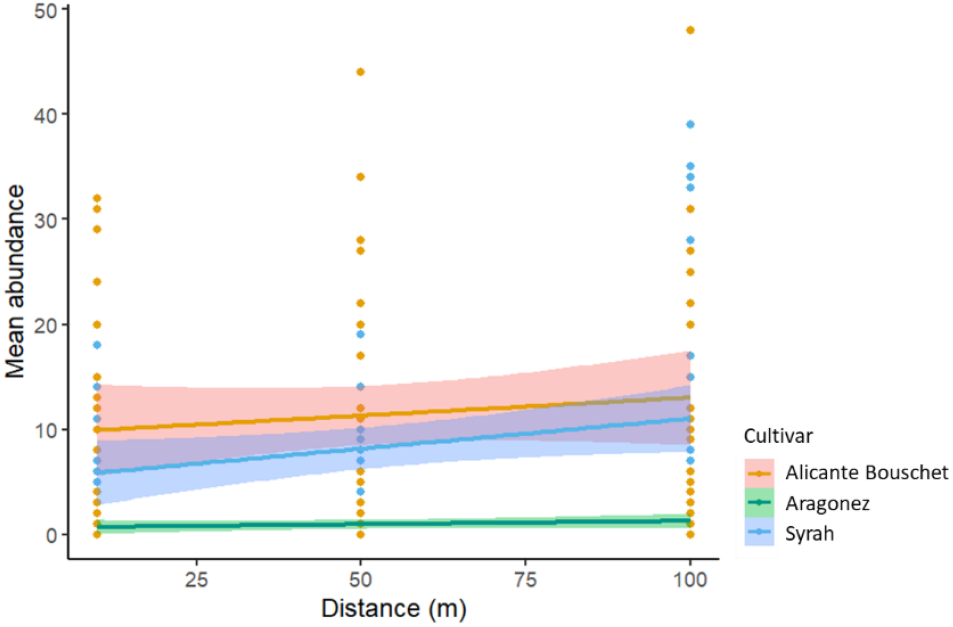
Mean abundance and 95% confidence interval of *Jacobiasca lybica* collected in 2022 in vineyards of each cultivar by distance to the periphery (m).

In 2022, the results in ‘Alicante Bouschet’ were very similarly to those of both years (2021 and 2022), with distance to the periphery (*χ*^2^ = 4.978, *df* = 1, *p* < 0.05) and periphery type (*χ*^2^ = 168.698, *df* = 1, *p* < 0.001) being both significant, but not their interaction (*χ*^2^ = 3.138, *df* = 1, *p* = 0.077) (Fig. 4a). However, in ‘Syrah’ the periphery type was significant (*χ*^2^ = 4.441, *df* = 1, *p* < 0.05) but not the distance to the periphery (*χ*^2^ = 3.728, *df* = 1, *p* = 0.054), and a significant interaction between distance and periphery was observed (*χ*^2^ = 17.824, *df* = 1, *p* < 0.001), with vineyards next to RH showing a downward slope in abundance per distance, whereas vineyards next to TH showed an upward slope (Fig. 4b). This means that distance to the periphery had opposite effects on *J. lybica* abundance in ‘Alicante Bouschet’ vs ‘Syrah’ vineyards next to RH (Fig. 4).

**Fig. 4.**
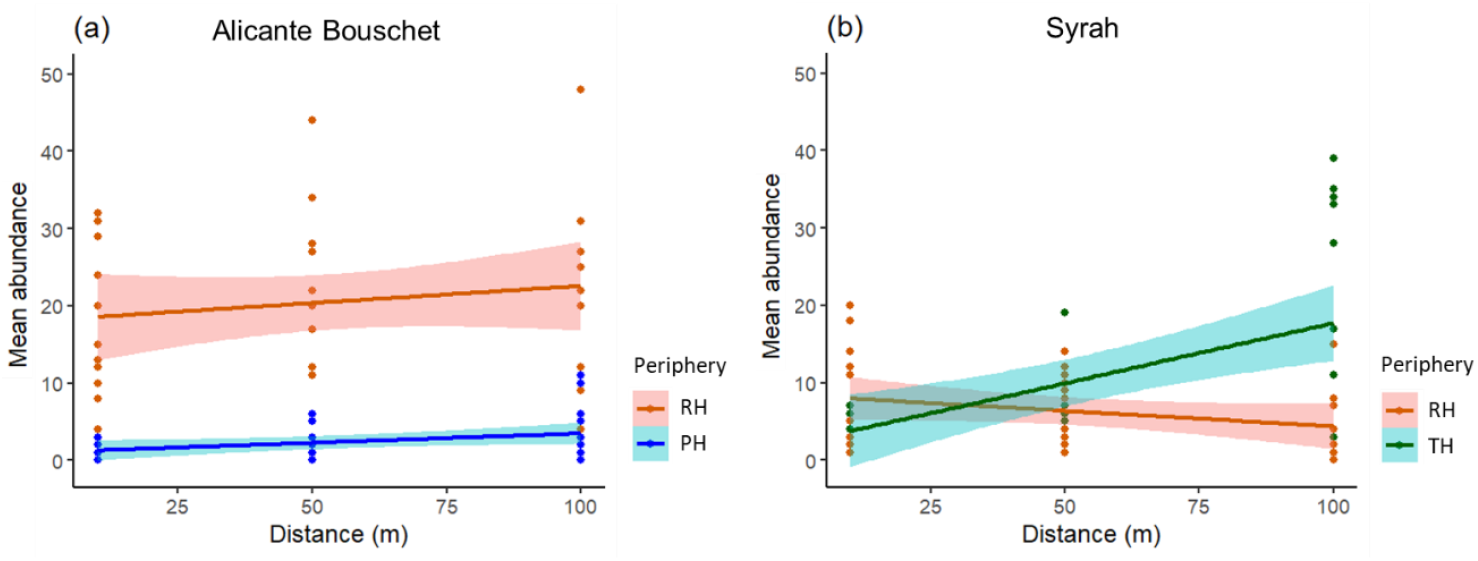
Mean abundance and 95% confidence interval of *Jacobiasca lybica* collected in 2022 in vineyards by distance to the periphery (m) and periphery type in: a) ‘Alicante Bouschet’; b) ‘Syrah’. PH – Planted Hedgerows; RH – Rosaceae-dominated Hedgerows; TH – *Tamarix*-dominated Hedgerows.

### Phytophagous mites

In peripheral plants, only eight taxa of phytophagous mites were collected. The most abundant species was the two-spotted spider mite, *T. urticae*, which was collected from two plant species: *F. angustifolia* and *R. canina*. Each of the other phytophagous mite taxa was collected from a single plant host, with samples from seven plant species not containing Tetranychidae or Tenuipalpidae phytophagous mites (Table 3).

**Table 3.**
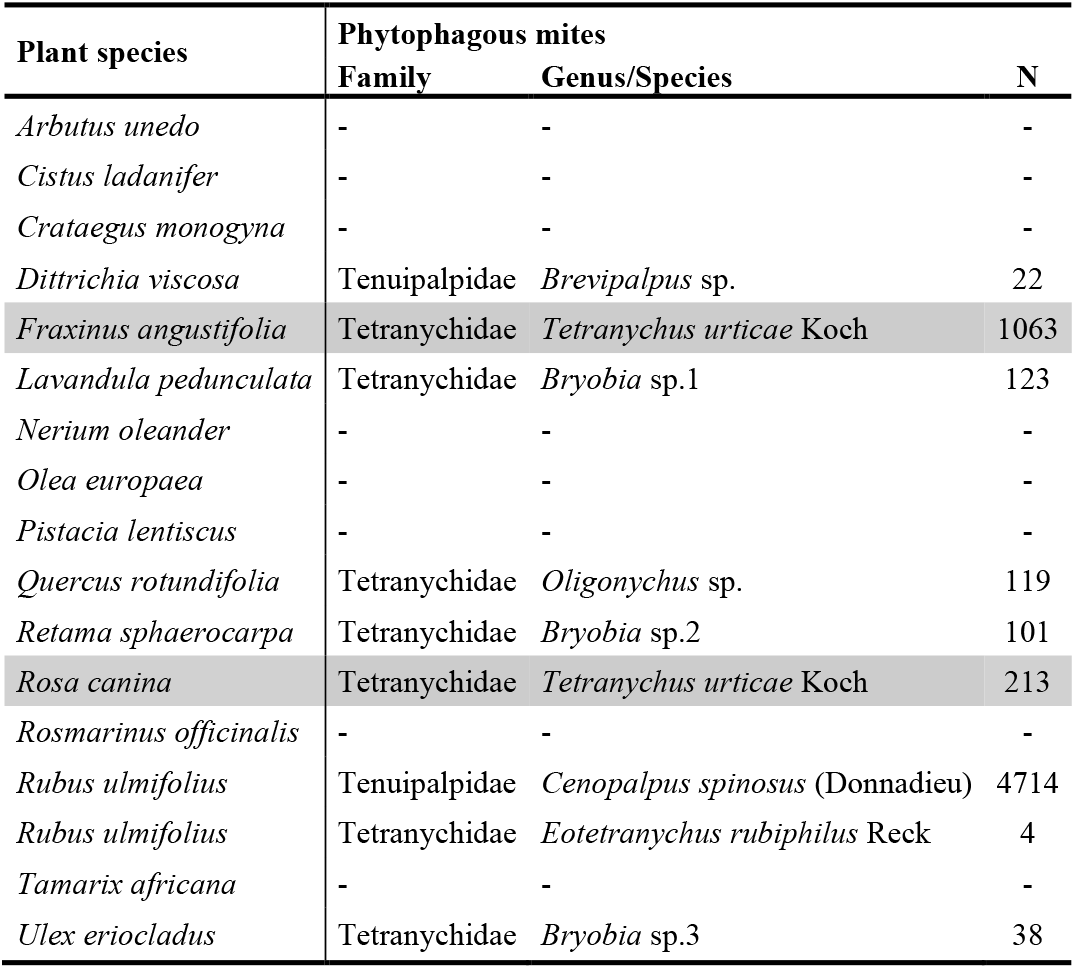
Presence and total abundance (N) of Tetranychidae and Tenuipalpidae phytophagous mites, by family and genus/species, collected during 2021 and 2022 in the sampled plant species. Gray lines indicate the presence of *Tetranychus urticae*.

In the vineyard, a total of 7245 mites were collected, of which 5734 (79.1%) were *T. urticae*, corresponding to 3947 specimens collected in 2021 and 1787 in 2022. This was the only spider mite collected from the vines, with the remaining mites collected belonging to the Families Acaridae, Phytoseiidae, Stigmaeidae, Tarsonemidae, Tydeidae, and to the Oribatida.

Distance to the periphery (*χ*^2^ = 24.458, *df* = 1, *p* < 0.001), plot type (*χ*^2^ = 28.271, *df* = 1, *p* < 0.001), cultivar (*χ*^2^ = 102.665, *df* = 3, *p* < 0.001), year (*χ*^2^ = 23.314, *df* = 1, *p* < 0.001), as well as most of their double and triple interactions significantly affected *T. urticae* abundance in the vineyard (Table S11).

In all cultivars, distance to the periphery (*χ*^2^ = 6.222, *df* = 1, *p* < 0.05), plot type (*χ*^2^ = 10.733, *df* = 1, *p* < 0.01), year (*χ*^2^ = 34.629, *df* = 1, *p* < 0.001), the interaction between plot type and year (*χ*^2^ = 8.28, *df* = 1, *p* < 0.01) and the interaction between distance, plot type and year (*χ*^2^ = 4.555, *df* = 1, *p* < 0.05) all significantly affected *T. urticae* abundance (Table S12).

Results were heterogeneous when looking at years separately: in 2021 only plot type was significant (*χ*^2^ = 15.555, *df* = 1, *p* < 0.001) due to a higher abundance of *T. urticae* in central plots (Fig. 5a), but distance (*χ*^2^ = 0.435, *df* = 1, *p* = 0.510) and interaction distance-plot type (*χ*^2^ = 0.113, *df* = 1, *p* = 0.737) were non-significant. Conversely, in 2022 plot types did not differ (*χ*^2^ = 0.058, *df* = 1, *p* = 0.809), but both distance (*χ*^2^ = 10.327, *df* = 1, *p* < 0.01) and distance-plot type (*χ*^2^ = 8.137, *df* = 1, *p* < 0.01) were significant, with increased distance leading to increased abundance of *T. urticae* only on peripheral plots (Fig. 5b).

**Fig. 5.**
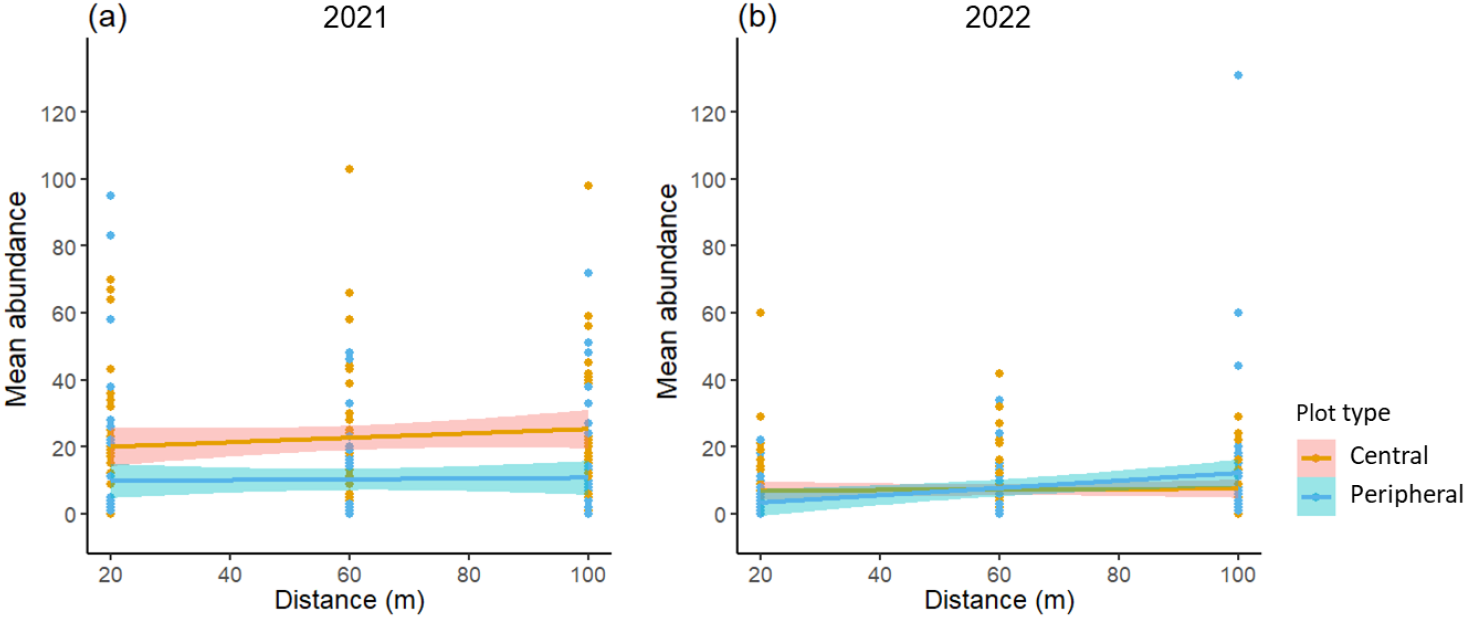
Mean abundance and 95% confidence interval of *Tetranychus urticae* collected in vine-yards of all cultivars by distance to the periphery (m), in Central and Peripheral plots: a) in 2021; b) in 2022.

Visual surveys indicated that spider mite attacks were, in general, relatively low and only exceeded economic threshold level in two central plots in ‘Syrah’ in 2020 and ‘Touriga Franca’ in 2022.

Analyzing data separately for each cultivar and year, distance to the periphery significantly affected the abundance of *T. urticae* in all cultivars except ‘Touriga Franca’, in at least one of the years (Table 4). However, in ‘Syrah’ there was a significant interaction between distance and plot type in both years (Table 4). Plot type also proved significant in most cultivars in both years (Table 4): although in most cases *T. urticae* was more abundant in central plots, peripheral plots harbored significantly more mites in ‘Alicante Bouschet’ in 2022 and in ‘Touriga Franca’ in 2021 (Fig. 6).

**Table 4.** Generalized linear model for *Tetranychus urticae* abundance (variable of interest) in each cultivar and year. *χ*^2^: chi-square value. *df*: degrees of freedom. ‘Distance’: distance to the periphery in each treatment (20 m, 60 m or 100 m). ‘Plot type’: location on the plot (in a central or peripheral position). AB - ‘Alicante Bouschet’; CS - ‘Cabernet Sauvignon’; Sy - ‘Syrah’; TF - ‘Touriga Franca’. ‘var’ – variable.

| Subset | Explanatory var. | $\chi^2$ | *df* | *p* value | Subset | Explanatory var. | $\chi^2$ | *df* | *p* value |
| --- | --- | --- | --- | --- | --- | --- | --- | --- | --- |
| in AB 2021 | Distance | 11.381 | 1 | <0.001 | in AB 2022 | Distance | 13.128 | 1 | <0.001 |
|  | Plot type | 12.686 | 1 | <0.001 |  | Plot type | 5.826 | 1 | <0.05 |
|  | Distance x Plot type | 0.117 | 1 | 0.733 |  | Distance x Plot type | 1.015 | 1 | 0.314 |
| in CS 2021 | Distance | 0.58 | 1 | 0.446 | in CS 2022 | Distance | 11.552 | 1 | <0.001 |
|  | Plot type | 42.202 | 1 | <0.001 |  | Plot type | 0.788 | 1 | 0.375 |
|  | Distance x Plot type | 3.122 | 1 | 0.077 |  | Distance x Plot type | 0.125 | 1 | 0.724 |
| in Sy 2021 | Distance | 7.313 | 1 | <0.01 | in Sy 2022 | Distance | 0.372 | 1 | 0.542 |
|  | Plot type | 96.273 | 1 | <0.001 |  | Plot type | 0.006 | 1 | 0.939 |
|  | Distance x Plot type | 5.03 | 1 | <0.05 |  | Distance x Plot type | 12.852 | 1 | <0.001 |
| in TF 2021 | Distance | 0.232 | 1 | 0.63 | in TF 2022 | Distance | 1.727 | 1 | 0.189 |
|  | Plot type | 8.455 | 1 | <0.01 |  | Plot type | 13.793 | 1 | <0.001 |
|  | Distance x Plot type | 0.002 | 1 | 0.969 |  | Distance x Plot type | 0.116 | 1 | 0.733 |

**Fig. 6.**
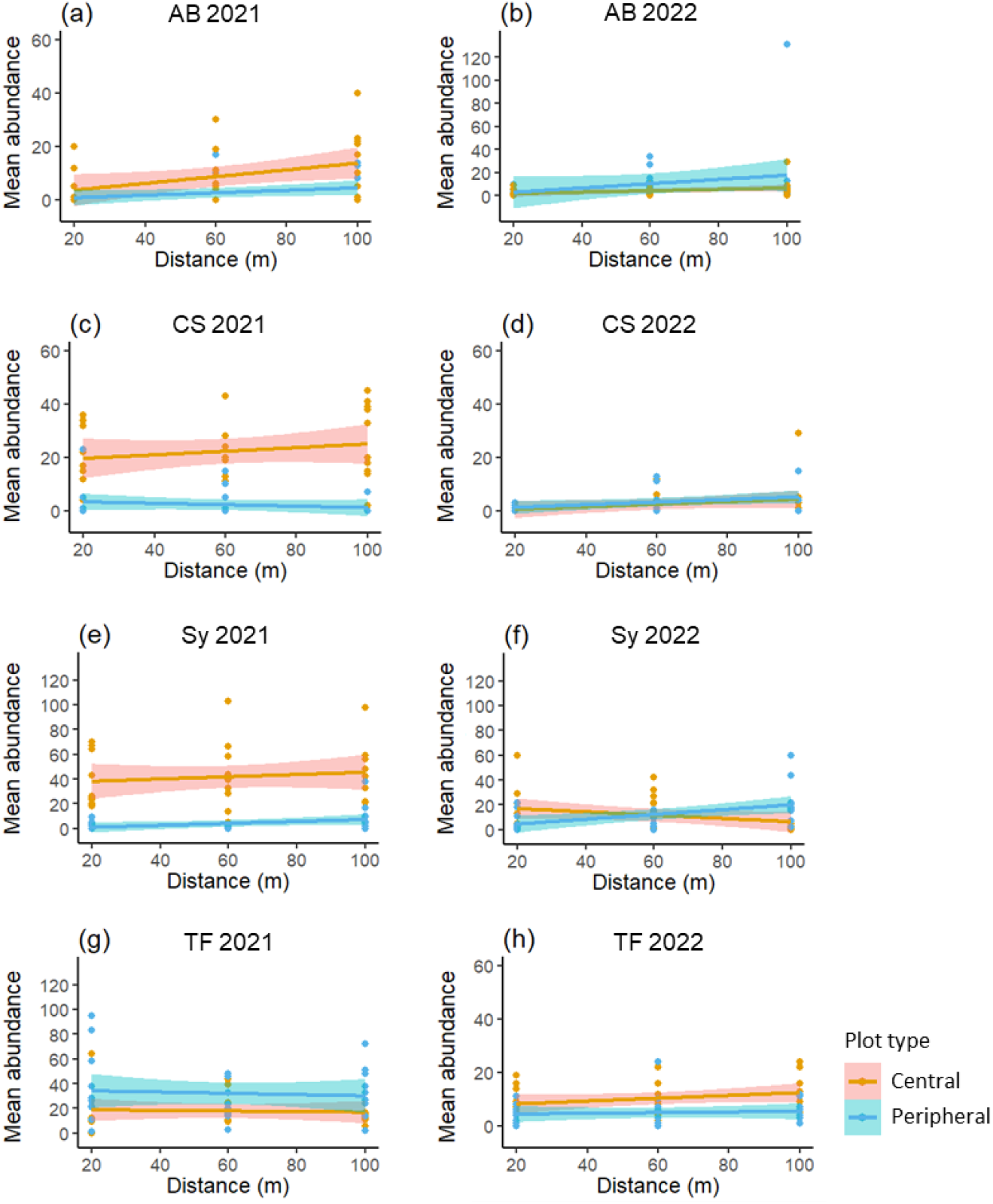
Mean abundance and 95% confidence interval of *Tetranychus urticae* collected in vineyards by distance to the periphery (m), in central and peripheral plots of: a) ‘Alicante Bouschet’ (AB) in 2021; b) ‘Alicante Bouschet’ (AB) in 2022; c) ‘Cabernet Sauvignon’ (CS) in 2021; d) ‘Cabernet Sauvignon’ (CS) in 2022; e) ‘Syrah’ (Sy) in 2021; f) ‘Syrah’ (Sy) in 2022; g) ‘Touriga Franca’ (TF) in 2021; h) ‘Touriga Franca’ (TF) in 2022. Note the differences in scales.

## Discussion

This study showed that, during the spring and summer months, the dominant pests in the sampled vineyards were *Jacobiasca lybica* and *Tetranychus urticae. Rubus ulmifolius* proved to be a winter repository of *J. lybica*, while *Rosa canina* and *Fraxinus angustifolia* hosted abundant populations of *T. urticae*. However, non-crop plants also harbored non-pest Auchenorrhyncha and mite species that might provide resources for natural enemies. In general, we detected a positive effect of plant-rich margins, with pest abundance in vineyards tending to decrease with proximity to the margin, and plots next to diverse peripheries having lower populations of *T. urticae* than plots surrounded by vineyards.

Some of the Cicadellidae detected in our study could not be identified to species level, and probably represent species which have not been formally described. This is the case for *Arboridia* sp.1, a new, yet undescribed species collected in the course of this work, associated with *R. ulmifolius*, a known host for the *Arboridia* genus (Ponti et al. 2005). Likewise, *Duilius* sp.1, collected in *Tamarix africana*, corresponds to a new species awaiting description. As with the leafhoppers, two of the *Bryobia* mite species (sp.2 and sp.3 in Table 3) most likely represent new, undescribed species, considering the distinctive morphological characters and the associated host plants. Furthermore, as far as we know, this is the first record of *Tamaricella tamaricis* (Puton) in Portugal, sampled as expected in *T. africana*. These new findings show that we are far from knowing in detail the Mediterranean arthropod fauna, its ecology and distribution, even in agroecosystems so intensively studied as vineyards, and that additional research is needed to fill the knowledge gaps.

The abundance of *J. lybica* was quite low in all peripheral perennial plants and ground cover vegetation. No plant was a significant repository of any Empoascini during summer, but *Rosmarinus officinalis* (in spring and winter) and *R. ulmifolius* (in winter) harbored the highest numbers of Empoascini and are, therefore, important repositories of this group. Importantly, the presence of *J. lybica* males in January in *R. ulmifolius*, albeit in low numbers, suggests that this plant is a winter host for this leafhopper. This may be contributing to an early buildup of its populations, resulting in higher abundance on adjacent vineyards in the summer. In fact, in August 2021 and 2022, ‘Alicante Bouschet’ vineyards next to Rosaceae-dominated Hedgerows (which contain *R. ulmifolius*) consistently showed significantly higher numbers of *J. lybica* than those next to planted hedgerows (that do not contain *R. ulmifolius*), and visual surveys showed that the former, but not the latter, regularly reached economic attack levels. Hence, despite *R. ulmifolius* known beneficial impact in biological control in vineyards (Altieri and Nicholls 2019), this species may represent some risk when next to vine cultivars particularly susceptible to *J. lybica*, such as ‘Alicante Bouschet’. Moreover, although *Hebata solani* was not statistically associated with *R. officinalis*, it is likely that this was due to a female-biased sex ratio in the population affecting the results of the statistical model, and that this plant is in fact a repository of *H. solani*. This hypothesis should be further studied, especially given the presence of this leafhopper on vineyard leaves in considerable numbers in the spring, a pattern previously observed in the study site (Alvarez 2020) and also in Italy (Mazzoni et al. 2001), and given the fact that *H. solani* is reported as a harmful pest in vineyards in Turkey (Alaserhat 2021).

*Rubus ulmifolius, Pistacia lentiscus* and *T. africana* proved to be repositories for other Auchenorrhyncha species. The association of *P. lentiscus* with *Bugraia ocularis* was known for the Mediterranean basin (D’Urso et al. 2019), as was the association of *T. africana* with *Opsius stactogalus* (D’Urso and Mifsud 2012) and *R. ulmifolius* with *Ribautiana tenerrima* (D’Urso et al. 2019; Ponti et al. 2003, 2005). The leafhopper *R. tenerrima* is a host for parasitoids of the genus *Anagrus* sp. (Hymenoptera, Mymaridae) (Altieri and Nicholls 2019; Ponti et al. 2003, 2005), known to be effective natural enemies of *J. lybica* (Hendawy et al. 2017; Klerks and van Lenteren 1991). As such, the high number of non-pest Auchenorrhyncha present in *Rubus* and *Tamarix* during spring and summer may serve as food resources for natural enemies (Ponti et al. 2005), resulting in an increase of their capacity to control pestiferous leafhoppers in the vineyards (Altieri and Nicholls 2019). Also, Rosaceae-dominated Hedgerows’ ground cover vegetation provided an important peak of Auchenorrhyncha in winter, indicating that ground cover herbaceous plants might also provide potential alternative resources for natural enemies without harboring harmful pests.

Concerning phytophagous mites, we identified two non-crop plant species hosting *T. urticae*, specifically *R. canina* and *F. angustifolia*, both known hosts of this species (Migeon and Dorkeld 2024), here reported for the first time as hosts in Western Europe. Other Tetranychidae and Tenuipalpidae phytophagous mites were detected on the plants surrounding the vineyards but, with the eventual exception of the *Brevipalpus* sp., all other phytophagous mites are monophagous or specialists on the genus/families of their plant host, and therefore do not represent a phytosanitary risk for the vineyards. In fact, their presence may be beneficial as they can be an alternative food source for generalist natural enemies.

*Nerium oleander* was the only plant present in the three sampled hedgerows, and it showed consistently low numbers of Auchenorrhyncha and no phytophagous mites, probably due to the high levels of toxicity which characterize this plant (Nia et al. 2018). Other species such as *Arbutus unedo* and *Olea europaea* also harbored low numbers of Auchenorrhyncha and no tetranychid or tenuipalpid mites. No plant species or ground cover proved to be a particularly good reservoir for *Xylella fastidiosa* potential vectors, which were collected in low numbers in both the periphery and vineyards.

The only phytophagous mite species detected in the vineyards, and usually below economic threshold levels, was the two-spotted spider mite *T. urticae*, a common pest that is wide-spread in Portugal associated with a wide range of crops, particularly fruit trees, tomato, strawberry, and vineyards (Naves et al. 2021; Zélé et al. 2018). In a similar study in South Africa, Vermaak (2019) detected eight species of tetranychid and tenuipalpid mites associated with vineyards, with *T. urticae* also being the dominant pest. In addition, other mites that can be important vine pests and cause conspicuous symptoms on the host, such as Eriophyidae (e.g., Duso et al. 2012), were not sampled in our study and their symptoms not observed in the sampled vineyards, suggesting their population levels were at most low. The reasons for the low pest incidence and damages are not clear, but this is an important observation in a property fully converted to certified organic farming since 2017, suggesting that organic practices can provide positive results regarding pest management.

In general, we found much higher numbers of pests on the vineyards than on peripheral habitats/species, which is in accordance with previous studies (Shapira et al. 2018). The abundance of both *J. lybica* and *T. urticae* in vineyards tended to correlate and increase with the distance to the periphery. However, in 2022 *T. urticae* abundance increased with distance to the periphery in peripheral plots but not in central plots (surrounded by other vineyards). This suggests that spider mite abundance increases with distance to marginal areas in particular when the latter are biodiverse. Abundance of pest species might therefore be dependent, at least partially, on marginal plant composition, with proximity to a biodiverse periphery being potentially beneficial in the reduction of pest numbers, as seen in previous studies (Altieri and Nicholls 2019; Favor et al. 2024; Gavinelli et al. 2020; Thomson and Hoffmann 2009, 2013).

Furthermore, central plots generally harbored more *T. urticae* than peripheral plots. Although we cannot establish causality, it is known that, at a landscape scale, the presence of seminatural habitats might be beneficial: Paredes et al. (2021) have shown that simplified vineyard-dominated landscapes have significantly higher pest outbreaks, when compared to complex landscapes with semi-natural habitats surrounding vineyards, and Dainese et al. (2017) showed that hedgerows did not increase biological control in a local scale in cereal adjacent fields, but did on a landscape scale.

The abundance of *T. urticae* showed considerable variations from year to year and with cultivar. Moreover, *J. lybica* populations were differentially affected by Rosacea-dominated hedgerows depending on the cultivar: increased distance to the hedgerows led to increasing pest numbers in ‘Alicante Bouschet’ but decreasing pest numbers in ‘Syrah’. A possible explanation lies in the fact that the hedgerow is in the middle of these two cultivars, hence *J. lybica* abundance in the two cultivars is changing in the same general direction, suggesting it is responding to a common environmental factor. Indeed, other important factors might be at play here, both for *T. urticae* and *J. lybica*, such as dominant wind direction responsible for the dispersal of arthropods (Altieri and Nicholls 2019; Duso et al. 2012; Tixier et al. 2006), management and composition of plant species of the ground cover between vineyard lines (Duso et al. 2012; Möth et al. 2021), terrain slope, humidity and temperature variations (Stavrinides et al. 2010) and/or higher exposition to the elements in peripheral vines, which we did not take into account. Moreover, the impact of the distance to the edge on pest crop abundance may hinge upon the relative impact of the margins on pests *vs* their natural enemies.

It is known that semi-natural areas around vineyards that provide non-crop plant resources, such as hedgerows and woody margins, can improve ecosystem connectivity (Altieri and Nicholls 2019; Boinot et al. 2023; Dainese et al. 2017; Garratt et al. 2017; Holland 2019), decrease pest outbreaks and pesticide use (Courson et al. 2024; Favor et al. 2024; Paredes et al. 2021). Our results underline the importance and potential benefits of biodiverse peripheral areas and the relevance of a good selection of plant species when managing or creating these areas. By unraveling the associations between phytophagous species and plant hosts, we hope to contribute with knowledge that allows for better management of such areas on the periphery of vineyards.

## Acknowledgments

We want to thank: Francesco Poggi and Marco de Haas for helping with Auchenorrhyncha identification; Ricardo Ramirez, Rui Félix, Sandra Antunes, Cândida Ramos, Mariana Azeredo, Emma Lopez and Carolina Matias for helping with data collection or sample processing; and José Santos for helping with the adaptation of suction equipment. We also thank the Herdade do Esporão agricultural team, in particular Amandio Rodrigues, Rui Flores and Carina Neto, for providing us total access to the property, support with travel and logistics for field work and access to data about agricultural practices.

## Funding

Renata Santos work was funded by the Foundation for Science and Technology (FCT), through the grant SFRH/BD/146090/2019. Herdade do Esporão provided funds for field work, field and lab materials. Authors received support from research centers: Linking Landscape, Environment, Agriculture and Food (LEAF) (https://doi.org/10.54499/UIDB/04129/2020) and Associate Laboratory TERRA (LA/P/0092/2020); Centre for Ecology, Evolution and Environmental Changes (cE3c) (https://sciproj.ptcris.pt/157405UID, https://doi.org/10.54499/UIDB/00329/2020, https://doi.org/10.54499/UIDP/00329/2020 and https://doi.org/10.54499/LA/P/0121/2020), and Centre for Environmental and Marine Studies (CESAM-Ciências) (https://doi.org/10.54499/UIDB/50017/2020, https://doi.org/10.54499/UIDP/50017/2020 and https://doi.org/10.54499/LA/P/0094/2020).

Sara Magalhães would like to acknowledge funding from HotPest: ref - 2022.04172.PTDC (https://doi.org/10.54499/2022.04172.PTDC); Pedro Naves from FCT the GREEN-IT Bioresources for Sustainability Unit (https://doi.org/10.54499/UIDB/04551/2020 and https://doi.org/10.54499/UIDP/04551/2020); and Leonor Rodrigues from a CEEC: ref - 2022.00518.CEECIND (https://doi.org/10.54499/2022.00518.CEECIND/CP1715/CT0007).

## Conflict of interest

The authors have no conflicts of interest.

## Notes

### Competing Interest Statement

The authors have declared no competing interest.

