## Supplementary material for "The association of Mediterranean plant species with herbivorous arthropods and its effect on pest abundance in organic vineyards": Table S2-S5-S6-S7-S8-S10-S11-S12

**Table S2** Generalized linear model output for Empoascini and other Auchenorrhyncha abundance. *χ*^2^ provides the chi-square value obtained in each analysis. *df* indicates the degrees of freedom. ‘Plant species’, refers to plants in all hedgerows (*Arbutus unedo*, *Nerium oleander* - tested separately for each hedgerow -, *Olea europaea*, *Pistacia lentiscus*, *Rosa canina*, *Rosmarinus officinalis*, *Rubus ulmifolius*, *Tamarix africana*); ‘Month’, month of sampling (May, August or January); ‘Year’, year of sampling (year 1 or year 2).

| **Variable of interest** | **Explanatory variable** | ***χ*^2^** | ***df*** | ***p* value** |
| --- | --- | --- | --- | --- |
| Empoascini | Plant species | 145.662 | 9 | < 0.001 |
|  | Month | 25.980 | 2 | < 0.001 |
|  | Year | 5.063 | 1 | < 0.05 |
|  | Plant species x Month | 57.631 | 15 | < 0.001 |
|  | Plant species x Year | 3.704 | 9 | 0.930 |
|  | Month x Year | 5.817 | 2 | 0.055 |
|  | Plant species x Month x Year | 14.414 | 15 | 0.494 |
| Other Auchenorrhyncha | Plant species | 1251.71 | 9 | < 0.001 |
|  | Month | 30.91 | 2 | < 0.001 |
|  | Year | 0.00 | 1 | 0.987 |
|  | Plant species x Month | 45.00 | 15 | < 0.001 |
|  | Plant species x Year | 12.61 | 9 | 0.181 |
|  | Month x Year | 3.92 | 2 | 0.141 |
|  | Plant species x Month x Year | 14.62 | 15 | 0.479 |

**Table S5** Generalized linear model output for total Auchenorrhyncha abundance (variable of interest) in peripheral ground cover plants. *χ*^2^ provides the chi-square value obtained in each analysis. *df* indicates the degrees of freedom. ‘Hedgerow’, refers to different hedgerow types (Planted Hedgerows, Rosaceae-dominated Hedgerows or *Tamarix*-dominated Hedgerows); ‘Month’, month of sampling (May, August or January); ‘Year’, year of sampling (year 1 or year 2).

| **Explanatory variable** | ***χ*^2^** | ***df*** | ***p* value** |
| --- | --- | --- | --- |
| Hedgerow | 4.297 | 2 | 0.117 |
| Month | 84.807 | 2 | < 0.001 |
| Year | 7.433 | 1 | < 0.01 |
| Hedgerow x Month | 55.972 | 3 | < 0.001 |
| Hedgerow x Year | 5.627 | 2 | 0.060 |
| Month x Year | 4.491 | 2 | 0.106 |
| Hedgerow x Month x Year | 5.118 | 3 | 0.163 |

**Table S6** Multiple comparisons for the significant explanatory variables of the analysis of total Auchenorrhyncha abundance in ground cover. Contrasts with Bonferroni corrections were performed to interpret which conditions significantly differed from each other. ‘Hedgerow type’, one of three hedgerows sampled (Planted Hedgerows; Rosaceae-dominated Hedgerows; *Tamarix*-dominated Hedgerows; here represented as ‘PH’, ‘RH’, ‘TH’, respectively); ‘Month’, one of three months sampled (August, January and May; here represented as ‘Aug’, ‘Jan’ and ‘May’, respectively).

| **Significant explanatory variable** | **Comparison** | **Estimate** | ***SE*** | ***t*-ratio** | ***p* value** |
| --- | --- | --- | --- | --- | --- |
| Hedgerow type x Month | RH Aug versus TH Aug | -0.9550 | 0.523 | -1.825 | 0.1416 |
|  | RH Aug versus PH Aug | -1.7356 | 0.498 | -3.484 | <0.01 |
|  | TH Aug versus PH Aug | -0.7806 | 0.417 | -1.874 | 0.1416 |
|  | RH Jan versus TH Jan | NA | NA | NA | NA |
|  | RH Jan versus PH Jan | 2.5033 | 0.346 | 7.239 | <0.001 |
|  | TH Jan versus PH Jan | NA | NA | NA | NA |
|  | RH May versus TH May | -0.2124 | 0.349 | -0.608 | 0.7533 |
|  | RH May versus PH May | 0.0957 | 0.359 | 0.266 | 1.000 |
|  | TH May versus PH May | 0.3081 | 0.356 | 0.865 | 0.6364 |
|  | RH Aug versus RH Jan | -4.6365 | 0.477 | -9.723 | <0.001 |
|  | RH Aug versus RH May | -2.5151 | 0.486 | -5.173 | <0.001 |
|  | RH Jan versus RH May | 2.1214 | 0.339 | 6.260 | <0.001 |
|  | TH Aug versus TH Jan | NA | NA | NA | NA |
|  | TH Aug versus TH May | -1.7726 | 0.400 | -4.437 | <0.001 |
|  | TH Jan versus TH May | NA | NA | NA | NA |
|  | PH Aug versus PH Jan | -0.3976 | 0.375 | -1.061 | 0.5236 |
|  | PH Aug versus PH May | -0.6838 | 0.375 | -1.822 | 0.1416 |
|  | PH Jan versus PH May | 0.2862 | 0.366 | 0.783 | 0.6528 |

**Table S7** Generalized linear model output for abundance of males of the most abundant Auchenorrhyncha species (global abundance of N≥1% of individuals) collected in the ground cover. *χ*^2^ provides the chi-square value obtained in each analysis. *df* indicates the degrees of freedom. ‘Hedgerow type’, type of hedgerow sampled (PH, RH or TH).

| **Variable of interest** | **Explanatory variable** | ***χ*^2^** | ***df*** | ***p* value** |
| --- | --- | --- | --- | --- |
| *Duilius* sp.1 | Hedgerow type | 6.5657 | 2 | < 0.05 |
| *Euscelis lineolata* | Hedgerow type | 1.2649 | 2 | 0.5313 |
| *Euscelis* sp.1 | Hedgerow type | 0.7397 | 2 | 0.6908 |
| *Zyginidia scutellaris* | Hedgerow type | 1.0974 | 2 | 0.5777 |

**Table S8** Multiple comparisons for the significant explanatory variable in the analysis of hedgerow type for *Duilius* sp.1 abundance in ground cover. Contrasts with Bonferroni corrections were performed to interpret which conditions significantly differed from each other. ‘Hedgerow type’, one of three hedgerows sampled (Planted Hedgerows; Rosaceae-dominated Hedgerows; *Tamarix*-dominated Hedgerows; here represented as ‘PH’, ‘RH’, ‘TH’, respectively). Statistical significance is represented with asterisks.

| **Significant explanatory variable** | **Comparison** | **Estimate** | ***SE*** | ***t*-ratio** | ***p* value** |
| --- | --- | --- | --- | --- | --- |
| Hedgerow type | RH versus TH | -0.4455 | 0.200 | -2.227 | <0.05 |
|  | RH versus PH | 0.0206 | 0.204 | 0.101 | 0.9197 |
|  | TH versus PH | 0.4661 | 0.202 | 2.305 | <0.05 |

**Table S10** Generalized linear model output for *J. lybica* abundance in August 2021 and 2022 in ‘Alicante Bouschet’. *χ*^2^ provides the chi-square value obtained in each analysis. *df* indicates the degrees of freedom. ‘Distance’, distance in each treatment (10 m, 50 m or 100m); ‘Plot’, location on the plot (next to PH or RH); ‘Year’, year of sampling (2021 or 2022).

| **Explanatory variable** | ***χ*^2^** | ***df*** | ***p* value** |
| --- | --- | --- | --- |
| Distance | 9.435 | 1 | <0.01 |
| Plot | 265.944 | 1 | <0.001 |
| Year | 26.154 | 1 | <0.001 |
| Distance x Plot | 1.386 | 1 | 0.239 |
| Distance x Year | 0.001 | 1 | 0.979 |
| Plot x Year | 0.003 | 1 | 0.954 |
| Distance x Plot x Year | 1.279 | 1 | 0.258 |

**Table S11** Generalized linear model output for *T. urticae* abundance (variable of interest). *χ*^2^ provides the chi-square value obtained in each analysis. *df* indicates the degrees of freedom. ‘Distance’, distance in each treatment (20 m, 60 m or 100m); ‘Plot type’, location on the plot in a Central or Peripheral position; ‘Cultivar’, one of four cultivars sampled (‘Alicante Bouschet’, ‘Syrah’, ‘Cabernet Sauvignon’ and ‘Touriga Franca’); ‘Year’, year of sampling (2021 or 2022).

| **Explanatory variable** | ***χ*^2^** | ***df*** | ***p* value** |
| --- | --- | --- | --- |
| Distance | 24.458 | 1 | <0.001 |
| Plot type | 28.271 | 1 | <0.001 |
| Cultivar | 102.665 | 3 | <0.001 |
| Year | 23.314 | 1 | <0.001 |
| Distance x Plot type | 3.411 | 1 | 0.065 |
| Distance x Cultivar | 16.943 | 3 | <0.001 |
| Plot type x Cultivar | 21.502 | 3 | <0.001 |
| Distance x Year | 1.402 | 1 | 0.236 |
| Plot type x Year | 46.672 | 1 | <0.001 |
| Cultivar x Year | 34.001 | 3 | <0.001 |
| Distance x Plot type x Cultivar | 15.828 | 3 | <0.01 |
| Distance x Plot type x Year | 0.918 | 1 | 0.338 |
| Distance x Cultivar x Year | 17.703 | 3 | <0.001 |
| Plot type x Cultivar x Year | 65.375 | 3 | <0.001 |
| Distance x Plot type x Cultivar x Year | 2.118 | 3 | 0.548 |

**Table S12** Generalized linear model output for *T. urticae* abundance (variable of interest) in all cultivars. *χ*^2^ provides the chi-square value obtained in each analysis. *df* indicates the degrees of freedom. ‘Distance’, distance in each treatment (20 m, 60 m or 100m); ‘Plot type’, location on the plot (in a Central or Peripheral position); ‘Year’, year of sampling (2021 or 2022).

| **Explanatory variable** | ***χ*^2^** | ***df*** | ***p* value** |
| --- | --- | --- | --- |
| Distance | 6.222 | 1 | <0.05 |
| Plot type | 10.733 | 1 | <0.01 |
| Year | 34.629 | 1 | <0.001 |
| Distance x Plot type | 2.195 | 1 | 0.139 |
| Distance x Year | 2.817 | 1 | 0.093 |
| Plot type x Year | 8.28 | 1 | <0.01 |
| Distance x Plot type x Year | 4.555 | 1 | <0.05 |
